# The transcriptional landscape of the developing chick trigeminal ganglion

**DOI:** 10.1101/2024.07.20.604400

**Authors:** Carrie E. Leonard, Alec McIntosh, Johena Sanyal, Lisa A. Taneyhill

**Author notes:** Funding information: NIH R01DE024217, R21DE033786.

## Abstract

The trigeminal ganglion is a critical structure in the peripheral nervous system, responsible for transmitting sensations of touch, pain, and temperature from craniofacial regions to the brain. Trigeminal ganglion development depends upon intrinsic cellular programming as well as extrinsic signals exchanged by diverse cell populations. With its complex anatomy and dual cellular origin from cranial placodes and neural crest cells, the trigeminal ganglion offers a rich context for examining diverse biological processes, including cell migration, fate determination, adhesion, and axon guidance. Avian models have, so far, enabled key insights into craniofacial and peripheral nervous system development. Yet, the molecular mechanisms driving trigeminal ganglion formation and subsequent nerve growth remain elusive. In this study, we performed RNA-sequencing at multiple stages of chick trigeminal ganglion development and generated a novel transcriptomic dataset that has been curated to illustrate temporally dynamic gene expression patterns. This publicly available resource identifies major pathways involved in trigeminal gangliogenesis, particularly with respect to the condensation and maturation of placode-derived neurons, thus inviting new lines of research into the essential processes governing trigeminal ganglion development.

## Introduction

The trigeminal ganglion is responsible for relaying sensations like touch, pain, and temperature from the face and oral cavity to the central nervous system^1^. Trigeminal nerve dysfunction causes disabling conditions including migraine headache, orofacial pain, and chronic dry eye, whose etiologies remain poorly understood^2–4^. Hereditary disorders associated with cranial sensory deficits are also explained, in part, by altered trigeminal nerve development^5–7^. Despite clear relevance to human health, the genetic landscape and molecular mechanisms underlying trigeminal ganglion neurodevelopment are still poorly characterized.

Peripheral nervous system development begins early during embryogenesis, at stages that are difficult to access in mammals. Avian studies have historically offered critical insight into the cellular lineage and molecular processes that govern cranial gangliogenesis^8–10^. Primary sensory neurons reside in peripheral sensory ganglia which are largely derived from the neural crest, a transient, multipotent progenitor population that gives rise to diverse tissues during vertebrate embryogenesis^11^. Likewise, cranial ganglia contain neural crest-derived sensory neurons and glia while also receiving neuronal contributions from specialized ectodermal thickenings called placodes^12^. While most cranial ganglia contain either neural crest- or placode-derived neurons that are spatially distinct, the trigeminal ganglion uniquely comprises a mixed neuronal population arising from both lineages^10,13^. Development of a functional trigeminal ganglion relies upon intrinsic transcriptional programming as well as reciprocal communication between neural crest and placode cells^14–17^.

Cranial neural crest cells, which originate in the dorsal neural folds and migrate to their final destinations within the embryo, influence the differentiation, migration, and growth of placode cells^18–20^. Chemotactic cues are sent by both neural crest and placode cells to direct their migration to the trigeminal ganglion anlage^21,22^. In tandem, migratory neural crest cells create corridors through which placodal neurons travel after delaminating from the overlying ectoderm^23,24^. Neurogenesis is staggered between the two progenitor populations, such that placode cells are the first to adopt a neuronal fate, followed by neural crest cells, which also give rise to glia, including satellite glial cells within the ganglion and Schwann cells that line the nerves^25,26^. The coalescence and condensation of placodal neurons with undifferentiated neural crest cells to form the ganglion is mediated by cell adhesion molecules including Cadherins^27,28^, and signaling pathways, like Wnt and Slit/Robo^28–30^. Once the earliest trigeminal sensory axons extend from placodal neurons, their growth toward craniofacial sites of innervation is further guided by diffusible factors, such as BMPs^31^, semaphorins^32,33^, and neurotrophins^34,35^. Thus, the developing trigeminal ganglion and its required interactions between neural crest cells and placodal neurons offer a rich context in which to investigate core biological processes, from fate determination to cell migration to axon guidance.

From decades of elegant work conducted in vertebrate embryos, there is a strong foundational understanding of neural crest and placode cell interactions and their contributions to cranial ganglia^36,37^. However, studies characterizing neurodevelopment from placodes are underrepresented compared to our knowledge of neural crest-derived neurons^38^. The early development and lack of genetic tools to target specific placodal subpopulations in mammalian models hinders research progress in this area. In contrast, chick embryos offer relative ease of access to facilitate the experimental manipulation of neural crest and distinct placodal progenitors^39^, with the additional benefit of continued development *in ovo*. With the evolutionary conservation of developmental mechanisms between species^37,40,41^, the chick embryo remains a favored, economical model for examining embryonic development, including the formation of neurogenic cranial placodes.

To elucidate normal gene expression dynamics during trigeminal ganglion development, we performed RNA-sequencing at sequential timepoints and curated a novel transcriptomic dataset to illustrate relative changes in gene expression over time. Raw and processed data are available in Gene Expression Omnibus (NCBI) under Accession #GSE282770 to encourage further exploration by researchers interested in craniofacial and/or peripheral nervous system development. Our findings identify key pathways involved in trigeminal gangliogenesis and provide critical insight into the molecular mechanisms underlying neurodevelopment and cellular interactions between placode-derived neurons and neural crest cells.

## Methods

### Collection of chicken embryos and trigeminal ganglia

Fertilized chicken eggs (*Gallus gallus*) were procured from Hy-Line North America (Elizabethtown, PA) or Centurion Poultry (Lexington, GA) and incubated in a humid environment at 37°C. Embryos were staged using the Hamburger-Hamilton (HH) method^42^, then removed from the egg and placed into sterile, ice-cold Ringer’s solution (0.125M sodium chloride, 1.5mM calcium chloride dihydrate, 5mM potassium chloride, 0.8mM sodium phosphate dibasic, adjusted to pH 7.4). Trigeminal ganglia were dissected using electrolytically sharpened tungsten needles, pooled by appropriate stages, and flash frozen in liquid nitrogen until total RNA extraction. With three pooled biological replicates for each stage, a total of 47 ganglia were used from HH13/14 embryos, 47 ganglia from HH15/16 embryos, and 56 ganglia from HH17/18 embryos. For immunohistochemical studies, HH17/18 embryos were collected and fixed with gentle agitation in 4% paraformaldehyde for 2 hours at room temperature, then washed 3 times for 10 minutes each in 1× PBS/0.1% Triton X-100 (Tx-100).

### RNA extraction, library preparation, RNA-sequencing and data processing

Total RNA was isolated according to the manufacturer’s protocol for the RNAqueous Total RNA Isolation kit (ThermoFisher, #AM1912). RNA concentration and integrity were quantified with Nanodrop (ThermoFisher) and Agilent 2100 bioanalyzer. The TruSeq sample preparation kit (Illumina) was used to generate poly(A)-enriched cDNA libraries, followed by assessment of quality and quantity using the bioanalyzer. 100bp paired-end reads were obtained with the Illumina HiSeq 1500 platform. Illumina adapter sequences were removed using Trimmomatic^43^ and the quality of reads assessed with FastQC^44^. HISAT2^45^ was used to align reads with a *Gallus gallus* reference genome (bGalGal1.mat.broiler.GRCg7b) and the number of reads per gene was quantified using HTSeq^46^. For principal component analysis, sample variability was calculated and visualized after normalization in R. Differential expression analyses were performed on normalized counts using DESeq2^47^, with three biological replicates from pooled trigeminal ganglia per indicated stage bin, and two technical replicates per biological replicate included. Heatmaps were generated using the online tool Morpheus from the Broad Institute (https://software.broadinstitute.org/morpheus/). Raw and processed data were deposited in Gene Expression Omnibus (NCBI) under Accession #GSE282770.

### Polar plot and Gene Ontology enrichment analyses

Normalized differential expression data were first used to calculate “H” by transforming sequential data (T^0^-T^1^-T^2^) into angular values representing relative gene expression at HH13/14 (T^0^), HH15/16 (T^1^), and HH17/18 (T^2^), respectively^48,49^. The determined angular values illustrate expression dynamics (i.e., low-low-high) on a radial plot. “H” corresponds to the relative angular position of individual gene expression across T^0^-T^1^-T^2^ (i.e., yellow; H=0 degrees indicates no change from T^0^-T^1^, and a significant increase in gene expression from T^1^-T^2^). MaxLog, as determined with DESeq2 as the maximal Log_2_ fold change between any two timepoints, is illustrated in the polar plot as distance from the center, while the transparency of a particular point reflects the average normalized gene expression of a given gene (i.e., values greater than 100 are fully opaque whereas lower averages are more transparent). Polar plots were generated with *ggplot2*^50^ and scaled expression profiles were created in R using previously described equations^51^ adapted from the Mandrup group^48,49^. Briefly, relative mean changes in gene expression were transformed to an angle between −180 and +180 degrees, then further transformed to a scale from 0 to 360 degrees, which was used to determine color based on degree. Differentially expressed genes were plotted using polar coordinates with the radius equal to MaxLog, and angular position around the radial plot determined by the change trend. Color-coded, three-point profile annotations demonstrating relative expression changes corresponding from left to right as HH13/14 (T^0^), HH15/16 (T^1^), and HH17/18 (T^2^), respectively, are displayed every 30 degrees along the outer edge of the polar plot. For Gene Ontology (GO) enrichment, genes were parsed into six sectors by grouping profiles into 60-degree intervals starting at 0 degrees, which denotes genes that were unchanged between T^0^-T^1^, but significantly increased from T^1^-T^2^. Using *EnrichR*^52–54^, genes within each 60-degree sector were assessed against the GO Biological Process and GO Cellular Compartment databases^55^. Prism (Graphpad) was used to graph the top fifteen pathways most enriched for each sector based on p-value. Excel files summarizing all color-matched calculated values for the polar plot and sector-parsed GO analyses are included as supplemental files.

### Embedding, cryosectioning, immunohistochemistry, and in situ hybridization data

Fixed embryos were processed through a sucrose gradient by submerging in 1× PBS plus 5% sucrose (w/v) for 10 minutes, followed by 1× PBS plus 15% sucrose (w/v) overnight at 4°C. Embryos were then incubated in 1× PBS plus 7.5% gelatin (w/v) and 15% sucrose at 37°C for 3 hours, before transfer into a solution of 1× PBS plus 20% gelatin (w/v) and 15% sucrose and incubation overnight at 37°C in a humidified incubator. Gelatin-equilibrated embryos were then embedded in 1× PBS plus 20% gelatin and flash frozen in liquid nitrogen vapor. Embedded embryos were sectioned at 12μm on a cryostat at −25°C, and tissue sections were collected on Superfrost Plus charged slides (VWR, Cat. No. 48311-703). Slides were de-gelatinized in 1× PBS at 42°C for 15 minutes, then rinsed in 1× PBS at room temperature before permeabilization in 1× PBS/0.5% Tx-100 for 10 minutes. Tissue was blocked with 10% (v/v) heat-treated sheep serum (HTSS, Sigma) in 1×PBS/0.1% Tx-100 for 1 hour at room temperature in a humidified chamber. Primary antibodies were diluted in 1× PBS/0.1% Tx-100 plus 5% HTSS and applied overnight at 4°C using the following concentrations: Sox10 (1:100, Cat. No. 66786-1-Ig, Proteintech); Islet1 (1:100, Cat. No. 40.2D6, DSHB); STMN2 (1:100, Cat. No. 720178, ThermoFisher); beta III Tubulin (“Tubb3”, 1:500, Cat. No. AB78078, Abcam); N-Cadherin (1:100, Cat. No. AB18203, Abcam); and HuC/D (1:100, Cat. No. A-21271, ThermoFisher). The following day, tissue sections were washed 4 times for 30 minutes each at room temperature in 1× PBS/0.1% Tx-100. Fluorescently labeled secondary antibodies (ThermoFisher: goat anti-mouse IgG_1_ Cat. No. A-21125 or A-21240, goat anti-mouse IgG_2a_ Cat. No. A-21131, goat anti-mouse IgG_2b_ Cat. No. A-21145, goat anti-rabbit Cat No. A-11034 or A-21245; and Southern Biotech: goat anti-mouse IgG_2a_ Cat. No. 1080-31) were diluted at 1:500 in 1× PBS/0.1% Tx-100 plus 5% HTSS and applied overnight at 4°C in a humidified chamber. Slides were washed as described above for the primary antibodies, and coverslips mounted using DAPI Fluoromount-G Mounting Medium (Southern Biotech, Cat. No. 0100-20). Images were acquired and processed using a Zeiss LSM 800 confocal microscope with Zen Blue software (Zeiss). *In situ* hybridization data from the developing trigeminal ganglion in embryonic day (E)11.5 mouse was obtained from the Allen Developing Mouse Brain Atlas^56^.

## Results and Discussion

To elucidate gene expression dynamics and identify key molecular drivers of trigeminal ganglion development, we performed a bulk RNA-sequencing time course assay. Trigeminal ganglia were collected from chicken embryos, pooled into three Hamburger-Hamilton (HH) stage-dependent bins^42^ (HH13/14, HH15/16, or HH17/18), and separated into three biological replicates per bin before RNA extraction and sequencing. A total of 47 trigeminal ganglia were used from HH13/14 embryos, corresponding to a time when placodal neurons and neural crest cells begin to coalesce within the trigeminal ganglion anlage. HH15/16 trigeminal ganglia (n=47) are in the process of condensing into a compact structure, comprising a mixture of neural crest cells and placodal neurons that have initiated neurite extension. By HH17/18 (n=56), the trigeminal ganglion is densely populated with placodal neurons that are actively undergoing axon growth and guidance, comingled with neural crest cells that have not yet differentiated into neurons or glia^12,57,58^. Principal component analysis of normalized gene counts revealed samples generally clustered according to developmental stage, with HH13/14 trigeminal ganglia least similar to HH17/18 trigeminal ganglia, and HH15/16 trigeminal ganglia falling in between the distributions of HH13/14 and HH17/18 samples (Supplemental Figure 1A). Our goal was not only to detect differentially expressed genes at each stage, but to visualize the dynamic patterns spanning these three developmental timepoints. To this end, polar plots were created to summarize the relative relationships of individual genes from HH13/14 to HH15/16 to HH17/18, and cluster genes into groups based on similar temporal profiles (Figure 1)^48,49^.

**Figure 1.**
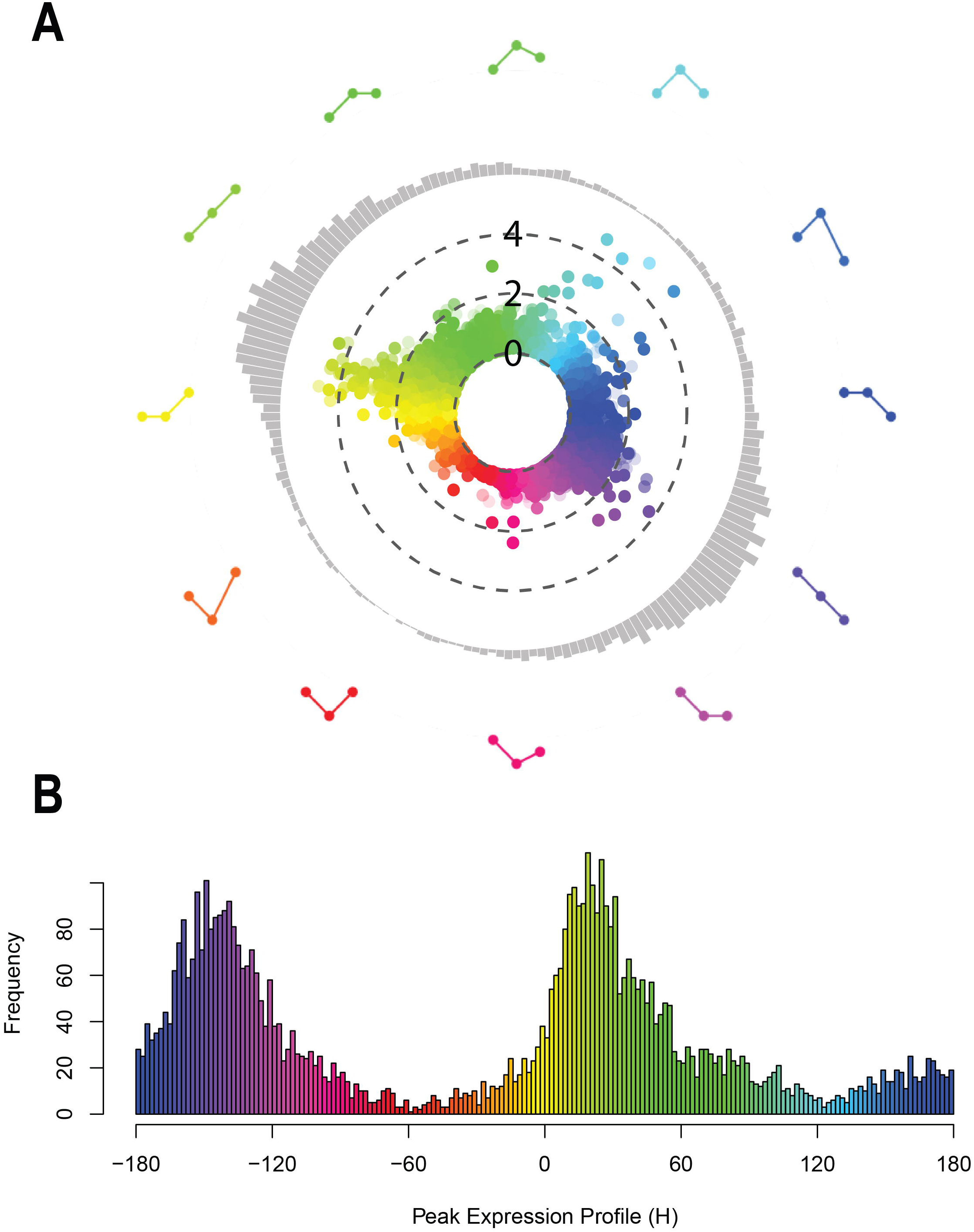
Temporally dynamic relative transcript expression during chick trigeminal ganglion development. **A)** Polar plot depicting relative temporal relationships of differentially expressed genes detected in the HH13/14, HH15/16, and HH17/18 chick trigeminal ganglion. Individual genes are represented as single points based on changes in expression across the three timepoints. In the three-point angular profiles surrounding the circular plot, the left dot represents HH13/14, the middle dot represents HH15/16, and the right dot represents HH17/18, whereas the pattern of each profile demonstrates the relative relationship of a gene’s expression at a given degree along the circular plot, i.e. the yellow profile represents a gene that does not change from HH13/14 to HH15/16, but is significantly upregulated from HH15/16 to HH17/18. Genes exhibiting a maximum Log_2_ fold change deemed significant (p<0.05 after Benjamini-Hochberg correction) at one timepoint relative to at least one other (three total comparisons) are shown. Maximal Log^2^ fold change for each gene is represented as distance from the center, while the concentric dotted lines represent Log_2_ fold change thresholds of 0, 2, and 4, respectively, from the center of the plot moving outward. Gray bars around the outer circumference demonstrate the number of genes at any given degree along the circle, which are transformed to a straight axis in B. **B)** Color-coded histogram illustrating the number of genes at any given degree (“H”) along the circumference of the polar plot, where 0 degrees equates to the yellow three-point profile in A.

In total, 14,506 genes were detected across the three timepoints. 5,703 genes exhibited significant differential expression (p<0.05 after Benjamini-Hochberg correction, see Supplemental File 1) at one timepoint compared to at least one other and were included in the polar plot (Figure 1A) and associated histogram (Figure 1B). The upregulation or downregulation of individual differentially expressed genes from HH13/14 to HH15/16 to HH17/18 was transformed into angular representations of relative expression over time, and placed onto a circular spectrum, where position around the circle is analogous to angular expression profiles, which are color-coded based on the degree within the circle, i.e., 0 degrees (yellow) is analogous to no change between HH13/14 and HH15/16, then a significant increase from HH15/16 to HH17/18 (Figure 1A). Of the differentially expressed genes, there were 765 with a maximal Log_2_ fold change (MaxLog) greater than 1.0, 124 genes with MaxLog greater than 2.0, and 14 genes with MaxLog exceeding 4.0 in any comparison, as indicated by increasing distance from the center of the polar plot (Figure 1A). Generally, the greatest changes in expression occurred in those genes exhibiting a temporal profile with upregulation between HH13/14 and HH15/16 (green to blue on polar plot), and either upregulation (yellow to green on polar plot) or sustained expression (green to blue on polar blot) from HH15/16 to HH17/18 (Figure 1A). The greatest numbers of differentially expressed genes occurred between 0 and +60 degrees (yellow to green) and between −120 and −180 degrees (indigo to purple) on the polar plot (Figure 1B), suggesting most changes in gene expression are unidirectional and/or sustained over time, regardless of increased versus decreased expression.

To understand how shared temporal gene expression profiles relate to biological events occurring in the developing trigeminal ganglion from HH13/14 to HH17/18, and to identify which genes may play a role in these processes, the polar plot was divided into 60-degree sectors. Differentially expressed genes within any given 60-degree sector displayed similar expression trajectories, on average, and were assessed for Gene Ontology (GO) enrichment against the GO: Biological Processes and GO: Cellular Component databases^55^ (Figures 2-4, Supplemental Figures 2-5, Supplemental File 2).

**Figure 2.**
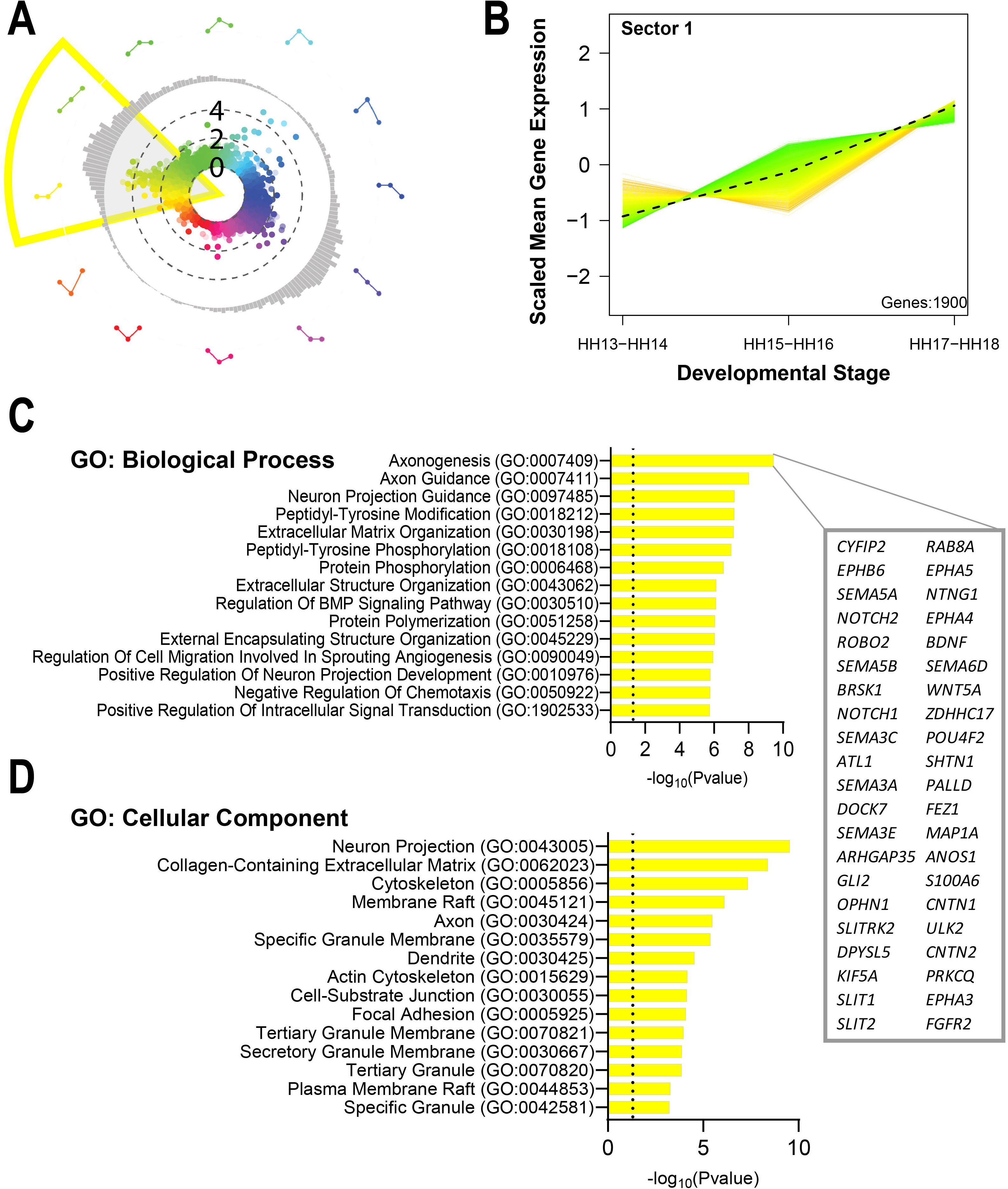
Genes associated with neuron development and axon guidance are continuously upregulated between HH13/14 and HH17/18 in the chick trigeminal ganglion. **A)** The yellow wedge overlayed on the polar plot represents the 60-degree angle used to define Sector 1 for gene ontology enrichment. **B)** Color-coded line plots reveal the relative expression patterns of individual genes included in Sector 1. The dotted line is the average scaled gene expression of all genes in Sector 1. **C-D)** Bar graphs illustrate the fifteen most enriched biological processes (C) and cellular components (D) associated with Sector 1 genes (Fisher’s exact test), in order of transformed p-value. The box outlined to the right lists all genes from Sector 1 associated with “Axonogenesis” (GO:0007409).

Within Sector 1, 1900 genes exhibited expression profiles that, on average, increased consistently from HH13/14 to HH15/16 and from HH15/16 to HH17/18, with some variation at the mid-point, but consistent upregulation by HH17/18 when compared to HH13/14 (Figure 2A and 2B). Significant biological processes and cellular components associated with Sector 1 genes included “Axonogenesis”, “Axon Guidance”, “Positive Regulation of Neuron Projection Development”, “Neuron Projection”, “Axon”, and “Cytoskeleton” (Figure 2C and 2D). Heatmaps illustrating relative temporal expression of differentially expressed genes associated with the regulation of axonogenesis revealed that, as expected, temporal expression patterns largely segregate according to sector. Further, the vast majority of genes associated with axonogenesis-related GO terms fall under Sector 1 (Supplemental Figure 1B). These findings align with the ongoing arrival and maturation of placodal neurons in the trigeminal ganglion between HH13/14 and HH17/18.

Indeed, we confirmed the presence of Sox10-positive neural crest cells and Islet1-positive placodal neurons in the HH18 trigeminal ganglion (Figure 3A). We also observed Sector 1 gene protein products, Stmn2 and Elavl4 (also known as HuD), in placodal neurons that expressed beta III tubulin or N-Cadherin (Figure 3B-C). Moreover, Sector 1 comprises several neuronal genes shown to be highly expressed in the developing mouse and chick trigeminal ganglion that play critical roles in axonal growth and maintenance, including *Ntrk1*, *Robo2*, and *Cntn2* (Figure 3D)^28,29,56,59–61^. While other signaling pathways known to be involved in sensory axon growth and guidance were among the upregulated genes in Sector 1, including Eph receptors, Semaphorins, and Notch (Figure 2C, right panel), genes within these families have not been well characterized in the trigeminal ganglion. Additionally, the GO enrichment results indicated changes in cell migration, cell signaling, and interactions with the extracellular matrix (Figure 2C and 2D). Such molecular changes may reflect migration dynamics of placodal neurons and/or neural crest cells as they populate the trigeminal ganglion and mature over time. They may also reflect genes important for sensing the environment as axons grow in search of target tissues, or as undifferentiated neural crest cells process extracellular cues that may influence their eventual fate.

**Figure 3.**
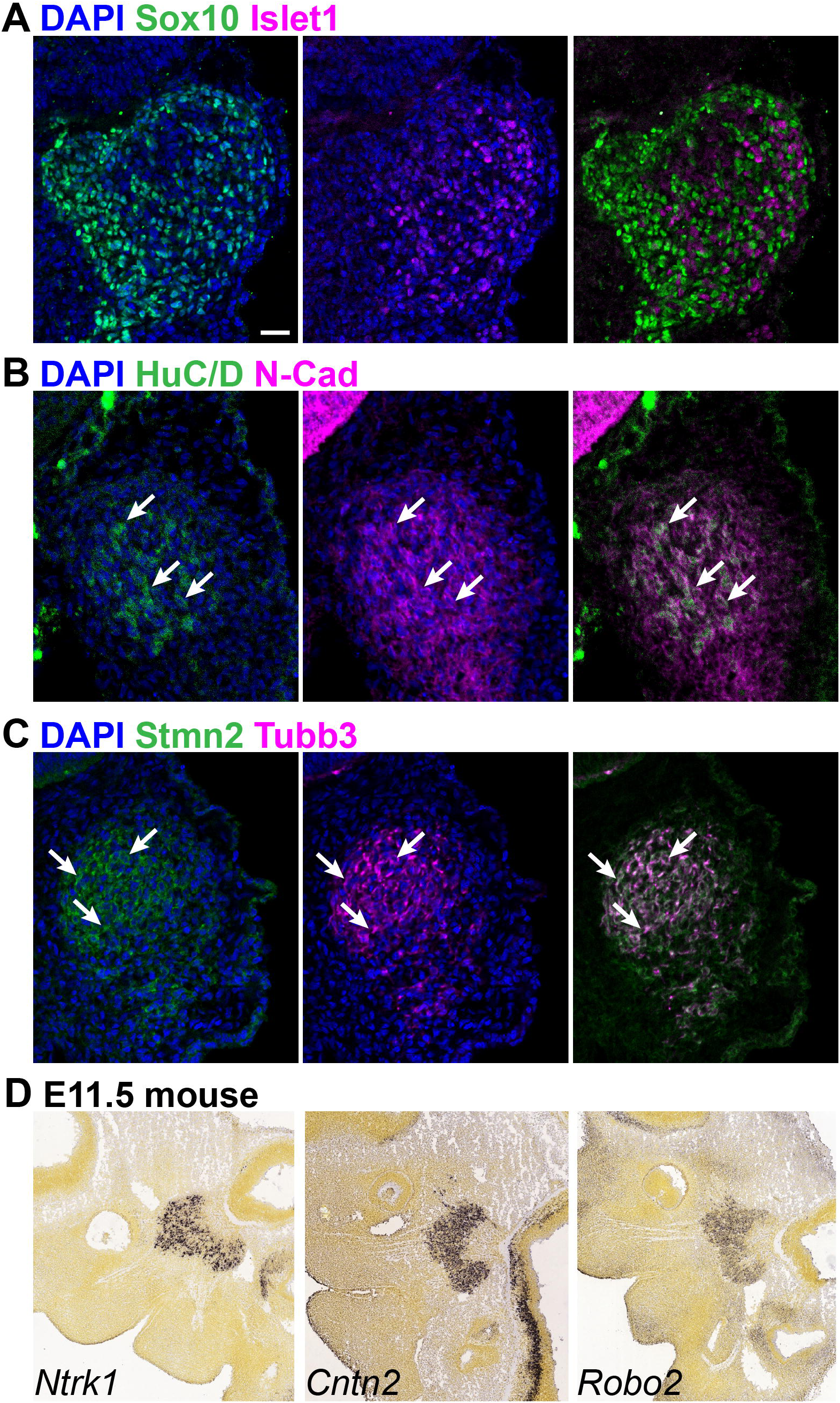
Validation of protein and transcript expression of neuronal genes in Sector 1. **A-C)** Fluorescent immunohistochemistry on transverse sections taken through the forming chick HH18 trigeminal ganglion demonstrates interspersed Sox10-positive (green) and Islet1-positive (purple) nuclei, confirming the presence of both neural crest cells and placodal neurons, respectively (A). Neurons expressing either N-Cadherin (B, purple) or beta III tubulin (“Tubb3”, C, purple) express protein products of genes in Sector 1, including HuD (B, green) and Stmn2 (C, green). Arrows point to representative cells that co-express N-Cadherin and HuD (B) or Tubb3 and Stmn2 (C). DAPI (blue) labels cell nuclei. **D)** *In situ* hybridization data from the Allen Developing Brain Atlas showing expression of Sector 1 genes in the E11.5 mouse trigeminal ganglion, including *Ntrk1* (https://developingmouse.brain-map.org/experiment/show/100046577), *Cntn2* (https://developingmouse.brain-map.org/experiment/show/100047095), and *Robo2* (https://developingmouse.brain-map.org/experiment/show/100046870)^56^. Scale bar in A is 20µm for A-C and 250µm for D.

In Sector 2, 796 genes, on average, were upregulated at HH15/16 compared to HH13/14, with sustained, but not further increased, expression at HH17/18 (Figure 4A and 4B). Differential expression of Sector 2 genes reflected activation of pathways involved in autophagy, as evidenced by enrichment in categories including “Lysosomal Transport”, “Positive Regulation of Autophagy”, “Vesicle-Mediated Transport”, “Lysosome”, “Recycling Endosome”, and “Autophagosome” (Figure 4C and 4D). While autophagy is a key player in processes including migration and differentiation that could be occurring in multiple cell types, it is also essential for proper neuronal development, plasticity, and survival^62,63^. Indeed, several neurodevelopmental disorders associated with sensory deficits, including autism spectrum disorders, are linked to dysfunctional autophagy^64^. Little has been described regarding the role of autophagy in trigeminal ganglion neurodevelopment, yet our findings indicate a large proportion of differentially expressed genes may participate in this process as placodal neurons mature.

**Figure 4.**
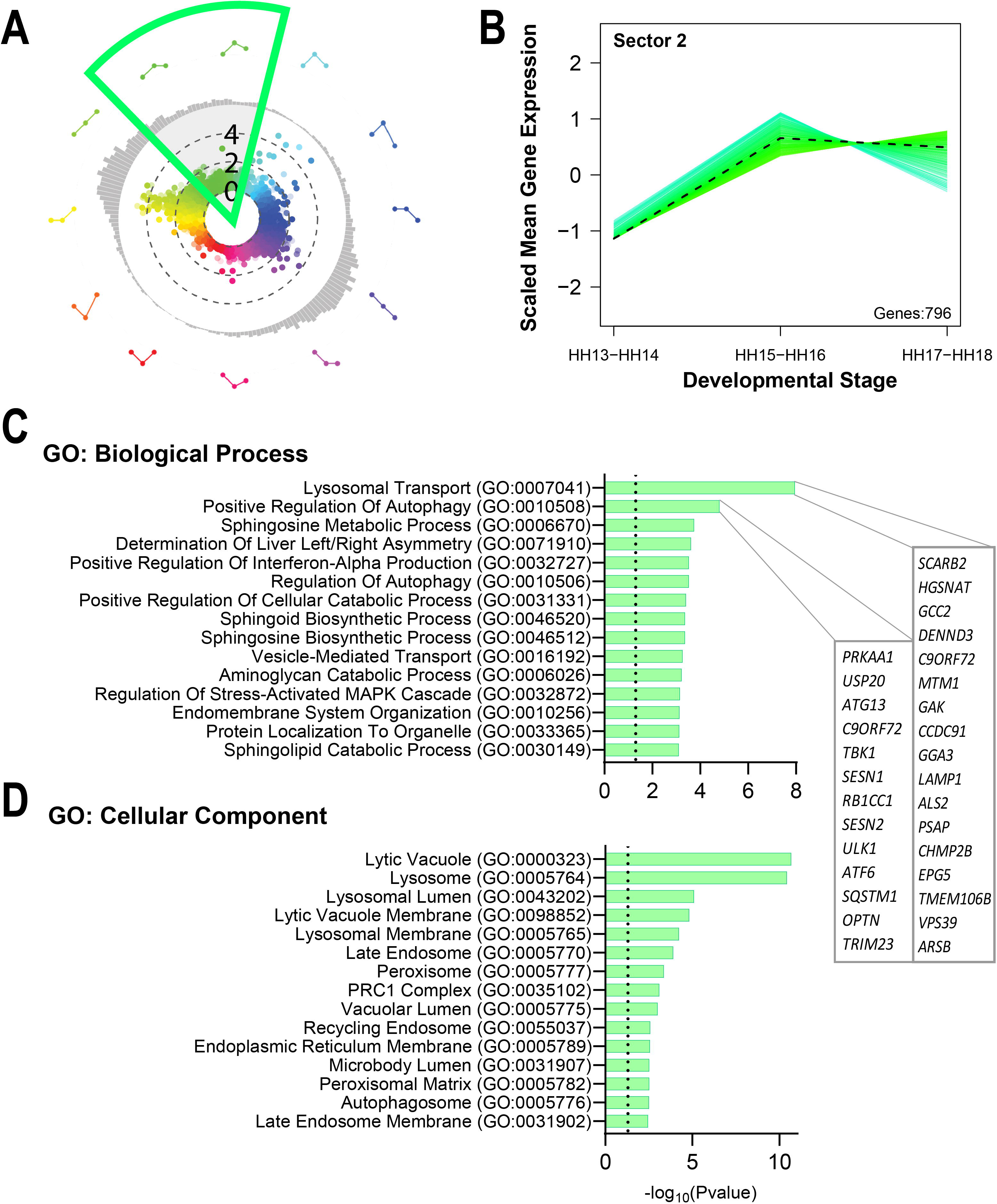
Autophagy-related genes are upregulated from HH13/14 to HH15/16, and maintained at HH17/18, in the chick trigeminal ganglion. **A)** The green wedge overlayed on the polar plot represents the 60-degree angle used to define Sector 2 for gene ontology enrichment. **B)** Color-coded line plots reveal the relative expression patterns of individual genes included in Sector 2. The dotted line is the average scaled gene expression of all genes in Sector 2. **C-D)** Bar graphs illustrate the fifteen most enriched biological processes (C) and cellular components (D) associated with Sector 2 genes (Fisher’s exact test), organized by transformed p-value. The box outlined to the right lists all genes from Sector 2 associated with “Lysosomal Transport” (GO:0007041) and “Positive Regulation of Autophagy” (GO:0010508).

While genes identified in Sectors 1 and 2 are largely associated with neurodevelopmental mechanisms, it is also plausible that they are driven by transcriptional changes in neural crest cells or even cranial mesenchyme, as multiple populations are present in our samples. Follow-up studies are needed to determine the unique cellular localization of differentially expressed transcripts, and how the identified pathways may function in placodal neurons versus neural crest cells at the examined timepoints.

Overall, genes in Sector 3 were associated with temporary upregulation of processes involved in sensory perception, glucose and ion transport, homeostasis, and cytoskeletal reorganization at HH15/16 (Supplemental Figure 2). Such processes are important for cell-cell communication, which is critical as undifferentiated neural crest cells and placodal neurons further condense to generate the ganglion, including supporting morphological changes in both cell populations^65–67^. Downregulated genes in Sectors 4 and 5 that remained lower at HH17/18 were generally associated with negative regulation of gene expression and protein biogenesis (Supplemental Figures 3-4). These findings may mirror trigeminal ganglion cells as they transition from proliferative and/or migratory progenitor states into cells with more defined fates, such as the increasing number of post-mitotic placodal neurons in the trigeminal ganglion over time, or neural crest cells that are preparing to differentiate into glia and/or neurons. Finally, the expression profiles of genes in Sector 6 generally reflected a temporary decrease in mitochondrial metabolic processes and cell motility (Supplemental Figure 5). This pattern may, too, reflect a pause and restructuring of energy-dependent cellular processes needed to support later steps in gangliogenesis, including axon extension and transport for neurodevelopment^68,69^. These non-exhaustive hypotheses require further investigation in order to fully understand the implications of individual gene expression trends in specific cell types.

Interpretation of our data is limited by the inability to assign changes in individual gene expression to a particular cell population. The RNA-sequencing data were generated from whole trigeminal ganglion lysate and, therefore, encompasses both placodal neurons and neural crest cells. Moreover, tissue samples likely included some contamination from the surrounding mesenchyme, especially at stages that precede trigeminal ganglion condensation. Given the migratory nature of several cell populations during the examined stages of craniofacial development, it is also possible that some transient changes in gene expression (for example, genes in Sectors 3 or 6) are explained by cells in transit to other tissues that do not ultimately remain in the trigeminal ganglion.

Even with these limitations, advantages of our bulk sequencing approach in comparison to single-cell or cell-sorted analyses include higher sensitivity and fewer gene “dropout events”, thus providing a more complete profile of average gene expression over time, which can be compared against more filtered data sets^70,71^. For example, Patthey et al. (2016) previously examined transcript expression in cranial sensory neurons by pre-selecting for mature neurons that expressed Neurofilament protein^58^. Our data confirmed the presence of neuronal genes deemed specific to the trigeminal ganglion, including *Prdm12*, *Gad2*, *Chrna3*, *Chrnb4*, and *Tox2* (Supplemental File 1)^58^. However, we also detected several genes in trigeminal ganglion lysate that Patthey et al. determined to be specific to neurons in other cranial ganglia, including *Esrrg*, *Irx2*, *Pdzrn4*, *Sult4a1*, *Shox2*, and *Lhx4* (Supplemental File 1)^58^. It is possible that these genes are instead expressed in non-neuronal cells within our trigeminal ganglion samples and, therefore, would have been selected against in Patthey et al. However, *Shox2* and *Lhx4*, for example, have established roles in sensory neuron development^72,73^. Moreover, our data capture genes that are expressed in newly differentiated, immature placodal neurons that would not yet express Neurofilament protein. Therefore, it is useful to consider filtered and bulk datasets together in order to draw and evaluate conclusions. In either case, RNA-sequencing, while informative, does not account for post-transcriptional or post-translational mechanisms that control gene expression. Follow-up assessment of protein products compared to their respective transcript profiles would provide additional insight into the functional consequences of the described transcriptional dynamics.

## Conclusions

Our report visually and quantitatively illustrates temporal dynamics in gene expression, cellular events, and key molecular pathways that participate in the development of the trigeminal ganglion in chick. These gene signatures correspond to key stages in trigeminal gangliogenesis, from the time when neural crest and placode cells coalesce and form a cohesive structure, through the neuronal maturation and early axon extension of placodal neurons. This novel, publicly available dataset is processed and curated with the goal of inspiring new lines of investigation into the molecular mechanisms underlying cranial sensory neuron development and the cellular interactions required for proper craniofacial development during embryogenesis.

## Acknowledgements

We thank Chyong-Yi Wu, Ph.D. for technical assistance in sample preparation. We thank Sue Zhao, Ph.D, Hector Corrada Bravo, Ph.D., and the UMD Institute for Bioscience & Biotechnology Research for their technical assistance and expertise in sequencing and preliminary data analyses.

## Supplemental Figures Legends

**Supplemental Figure 1.**
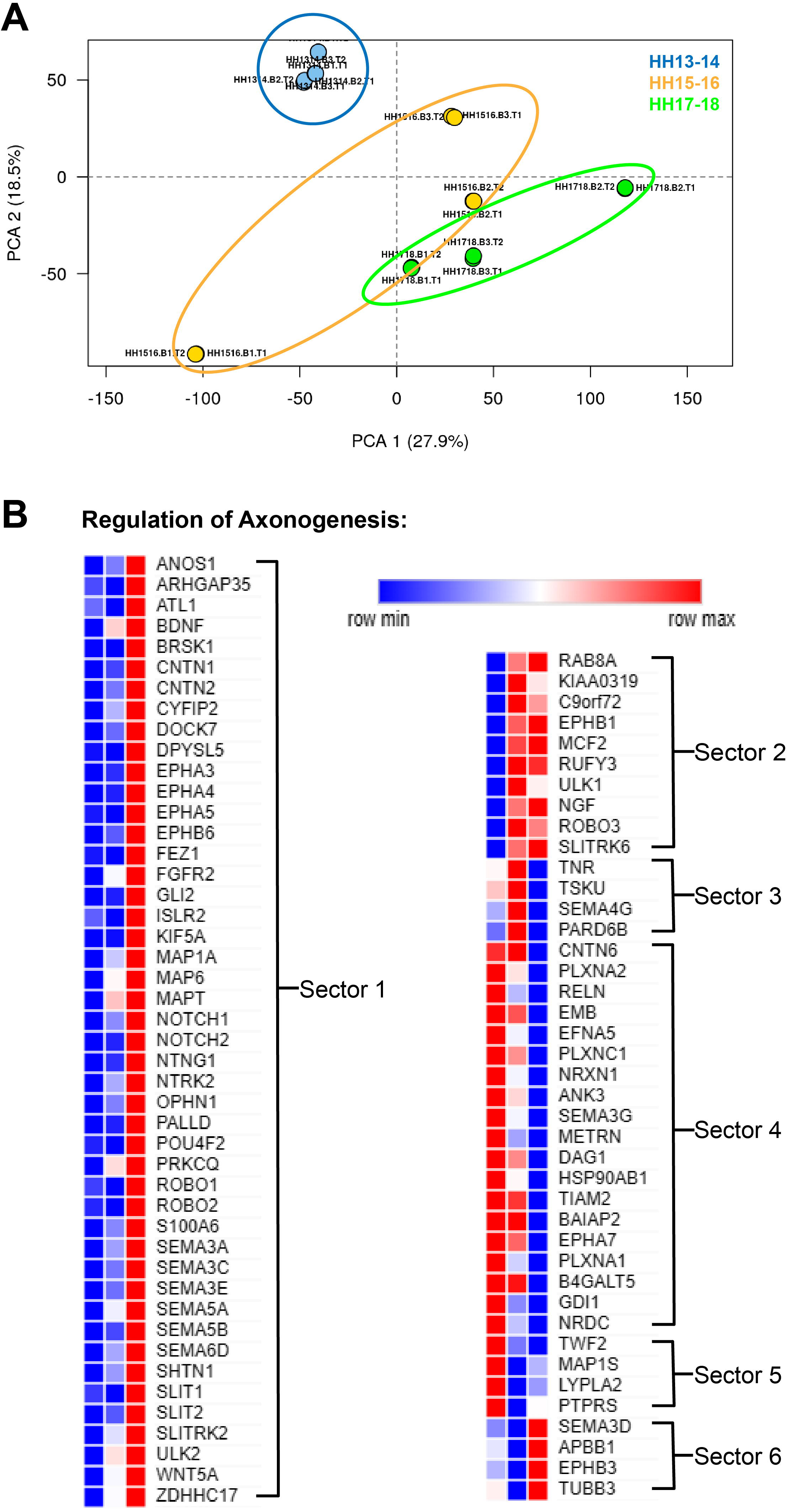
Analyses of sample similarity and temporal relationships of genes related to axonogenesis. **A)** Results of principal component analysis comparing all biological and technical replicates from HH13/14 (blue), HH15/16 (yellow), and HH17/18 (green) after normalization. Ellipses are overlayed to demonstrate how samples within each staging bin cluster together on the plot. **B)** Heatmap illustrating temporal expression patterns of genes associated with GO terms related to axonogenesis, including “Axonogenesis”, “Regulation of Axonogenesis”, “Positive/Negative Regulation of Axonogenesis”, “CNS Neuron Axonogenesis”, and “CNS Projection Neuron Axonogenesis”. Heatmap depicts relative expression at HH13/14 (left column), HH15/16 (middle column), and HH17/18 (right column), where blue represents the minimum expression for a given row, red represents the maximum expression, and white represents the mid-point between minimum and maximum expression. Brackets label the Polar Plot Sector under which each group of genes falls.

**Supplemental Figure 2.**
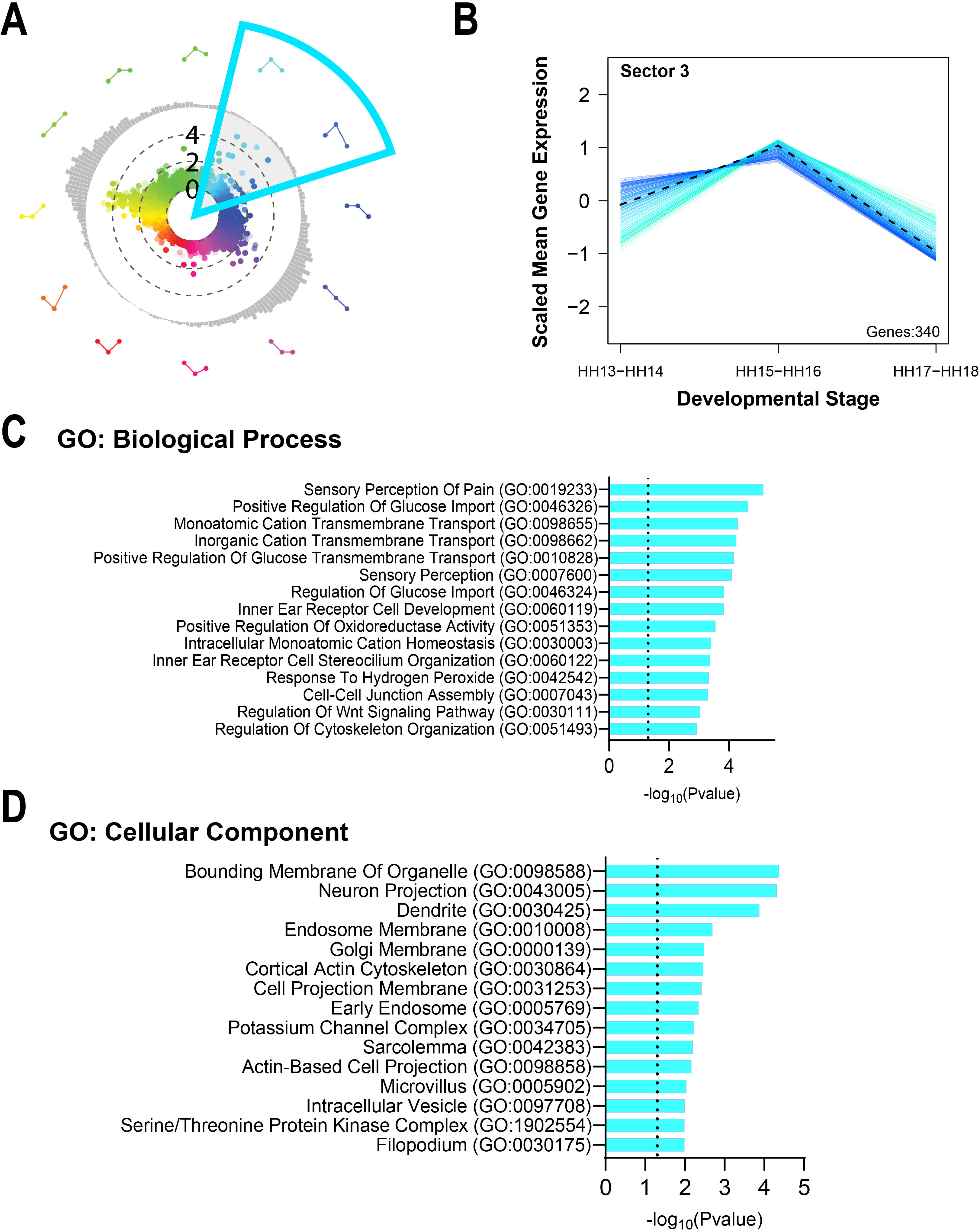
Sector 3 gene expression and ontologies. **A)** The light blue wedge overlayed on the polar plot from Figure 1 represents the 60-degree angle used to define Sector 3 for gene ontology enrichment. **B)** Color-coded line plots reveal the relative expression patterns of individual genes included in Sector 3. The dotted line is the average scaled gene expression of all genes in Sector 3. **C-D)** Bar graphs illustrate the fifteen most enriched biological processes (C) and cellular components (D) associated with Sector 3 genes (Fisher’s exact test), organized by transformed p-value.

**Supplemental Figure 3.**
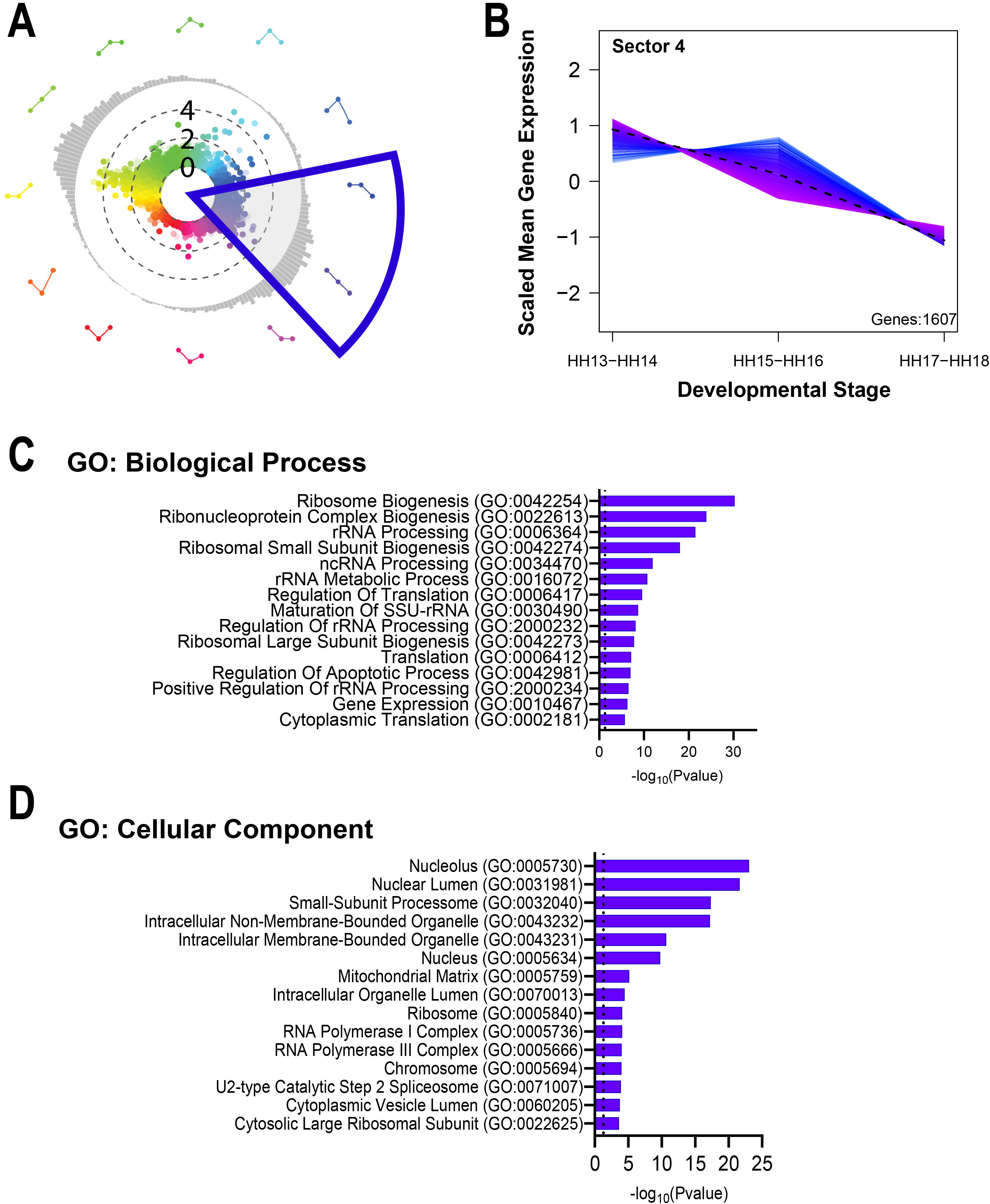
Sector 4 gene expression and ontologies. **A)** The dark blue wedge overlayed on the polar plot from Figure 1 represents the 60-degree angle used to define Sector 4 for gene ontology enrichment. **B)** Color-coded line plots reveal the relative expression patterns of individual genes included in Sector 4. The dotted line is the average scaled gene expression of all genes in Sector 4. **C-D)** Bar graphs illustrate the fifteen most enriched biological processes (C) and cellular components (D) associated with Sector 4 genes (Fisher’s exact test), organized by transformed p-value.

**Supplemental Figure 4.**
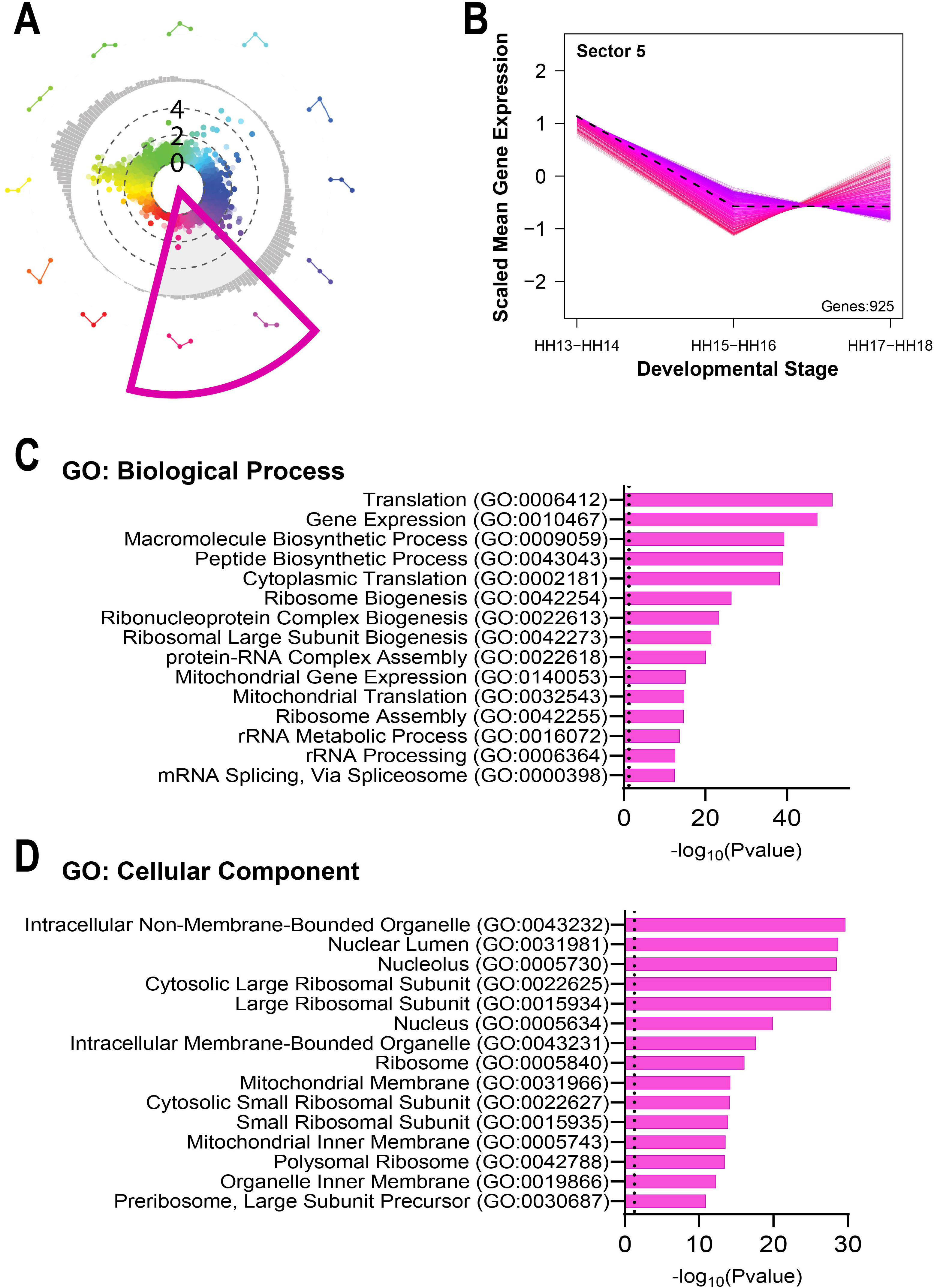
Sector 5 gene expression and ontologies. **A)** The pink wedge overlayed on the polar plot from Figure 1 represents the 60-degree angle used to define Sector 5 for gene ontology enrichment. **B)** Color-coded line plots reveal the relative expression patterns of individual genes included in Sector 5. The dotted line is the average scaled gene expression of all genes in Sector 5. **C-D)** Bar graphs illustrate the fifteen most enriched biological processes (C) and cellular components (D) associated with Sector 5 genes (Fisher’s exact test), organized by transformed p-value.

**Supplemental Figure 5.**
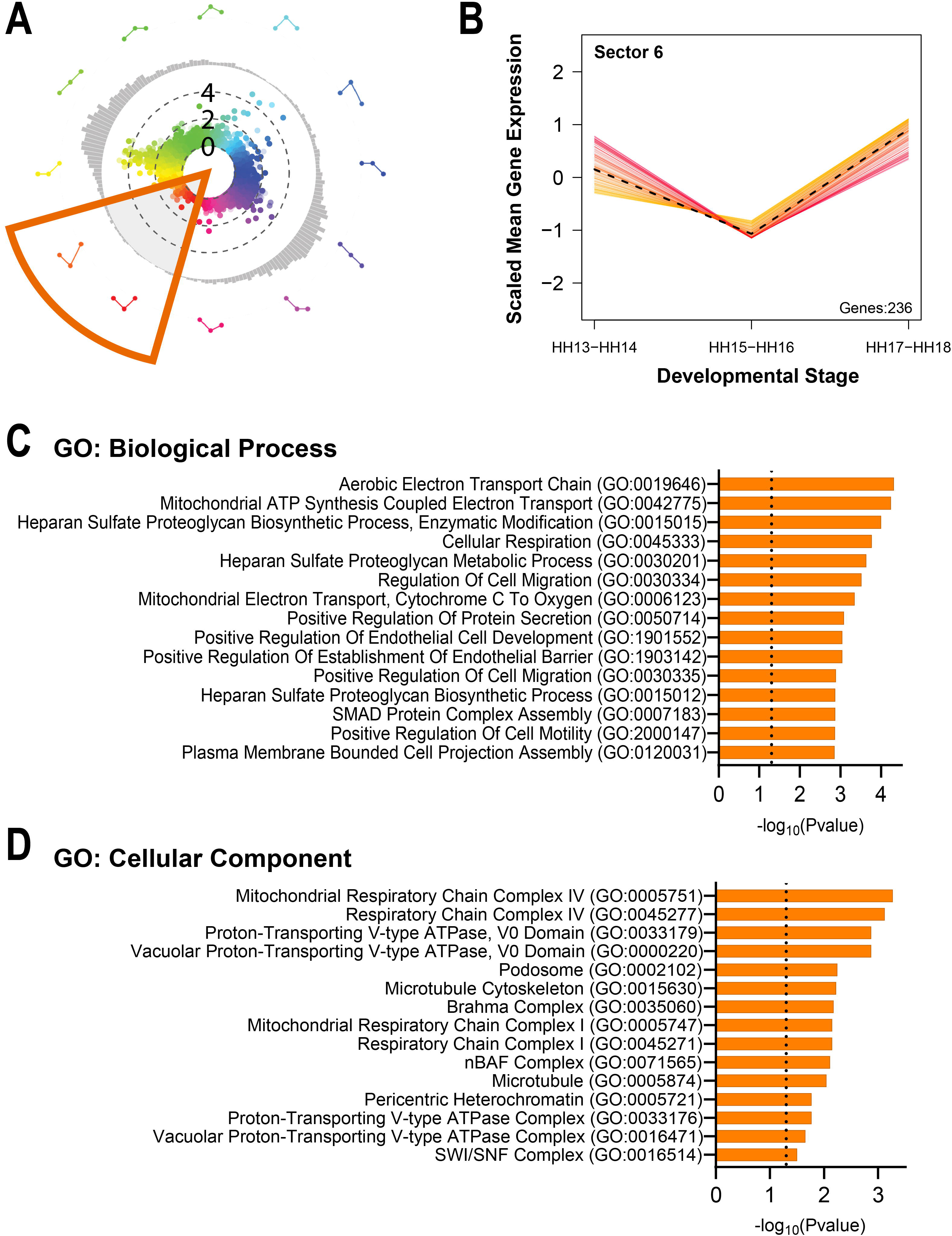
Sector 6 gene expression and ontologies. **A)** The orange wedge overlayed on the polar plot from Figure 1 represents the 60-degree angle used to define Sector 6 for gene ontology enrichment. **B)** Color-coded line plots reveal the relative expression patterns of individual genes included in Sector 6. The dotted line is the average scaled gene expression of all genes in Sector 6. **C-D)** Bar graphs illustrate the fifteen most enriched biological processes (C) and cellular components (D) associated with Sector 6 genes (Fisher’s exact test), organized by transformed p-value.

## Supplemental Data Files

**Supplemental File 1.** Color-coded Excel spreadsheet containing calculated values for differential expression comparisons and polar plot.

**Supplemental File 2.** Excel spreadsheets compiling gene ontology enrichment results for Biological Processes (BP) and Cellular Components (CC) associated with each 60-degree polar plot sector.

